# Evolutionary and homeostatic changes in morphology of visual dendrites of Mauthner cells in *Astyanax* blind cavefish

**DOI:** 10.1101/2020.05.13.094680

**Authors:** Zainab Tanvir, Daihana Rivera, Kristen E. Severi, Gal Haspel, Daphne Soares

**Author notes:** Corresponding Author, Daphne Soares, Biological Sciences, New Jersey Institute of Technology, 100 Summit street, Newark, NJ, 07102, USA.

## Abstract

Mauthner cells are the largest neurons in the hindbrain of teleost fish and most amphibians. Each cell has two major dendrites thought to receive segregated streams of sensory input: the lateral dendrite receives mechanosensory input while the ventral dendrite receives visual input. These inputs, which mediate escape responses to sudden stimuli, may be modulated by the availability of sensory information to the animal. To understand the impact of the absence of visual information on the morphologies of Mauthner cells during development and evolutionary time scales, we examined *Astyanax mexicanus*. This species of tetra is found in two morphs: a seeing surface fish and a blind cavefish. We compared the structure of Mauthner cells in surface fish raised under daily light conditions, surface fish that raised in constant darkness, and two independent lineages of cave populations. The length of ventral dendrites of Mauthner cells in dark-raised surface larvae were longer and more branched, while in both cave morphs the ventral dendrites were smaller or absent. The absence of visual input in surface fish with normal eye development leads to a homeostatic increase in dendrite size, whereas over evolution, the absence of light led to the loss of eyes and a phylogenetic reduction in dendrite size. Consequently, homeostatic mechanisms are under natural selection that provide adaptation to constant darkness.

## Introduction

Startle responses are rapid reactions that move an animal away from perceived danger (Peek and Card 2016). These behaviors are found in many species, and are typically mediated by fast-acting neural circuits characterized by large diameter neurons and reduced numbers of synapses (Eaton and Hackett 1984; Eaton et al., 1985). Mauthner cells are a pair of large neurons found in the hindbrain of teleost fishes that mediate escape responses (Bartelmez 1915; Eaton et al., 2001). These neurons receive input from sensory receptors and project contralaterally descending axons onto primary motor neurons in the spinal cord. Activation of a Mauthner cell results in rapid contraction of the contralateral musculature of the body trunk (Faber et al., 1989). These contractions generate a stereotypical behavior known as a C-start (Eaton et al., 1991).

Each Mauthner cell has two large-diameter primary dendrites. Each dendrite receives independent sensory information: acoustic and mechanosensory information on the lateral dendrite (Zottoli, 1977; Eaton et al., 1988) and visual and somatosensory information on the ventral dendrite (Faber and Korn, 1978; Zottoli et al., 1987, 1995; Canfield 2003). While much attention has been focused on inputs to the lateral dendrite, less is known about visual inputs to the ventral dendrite. Visual inputs mediated by the ventral dendrite reliably produce rapid escape responses across teleost species (Zottoli et. al 1987; Canfield 2006; Preuss et al., 2006; Dunn et al., 2016).

Troglobitic species throughout the world have convergent adaptations to life in caves, often including dramatic reductions of visual systems. Evolution in continual darkness has led to the degeneration of the visual system, and as animals become more cave-adapted, their eye structures are reduced and eventually vanish completely (Soares and Niemiller 2013; Barr, 1968; Fong et al., 1995; Protas and Jeffery, 2012; Culver and Pipan, 2016). The Mexican tetra fish, *Astyanax mexicanus* (Fig. 1), is currently extant in two morphs: an ancestral, river-dwelling morph (surface fish) and various derived, cave-dwelling morphs. Each lineage has unique adaptations to troglobitic life and is named after the cave in which they are endemic (Fig. 1A). There are two main lineages of fish that contain eyeless fish. The older lineage includes *Astyanax* cavefish from the Subterráneo, Pachón, and Chica populations (Dowling et al., 2002; Porter at al., 2007; Gross, 2012; Bradic et al., 2013). The younger lineage includes cavefish from the Los Sabinos, Curva, and Tinaja populations. These two populations are morphologically distinct (Dowling et al., 2002).

**Fig. 1.**
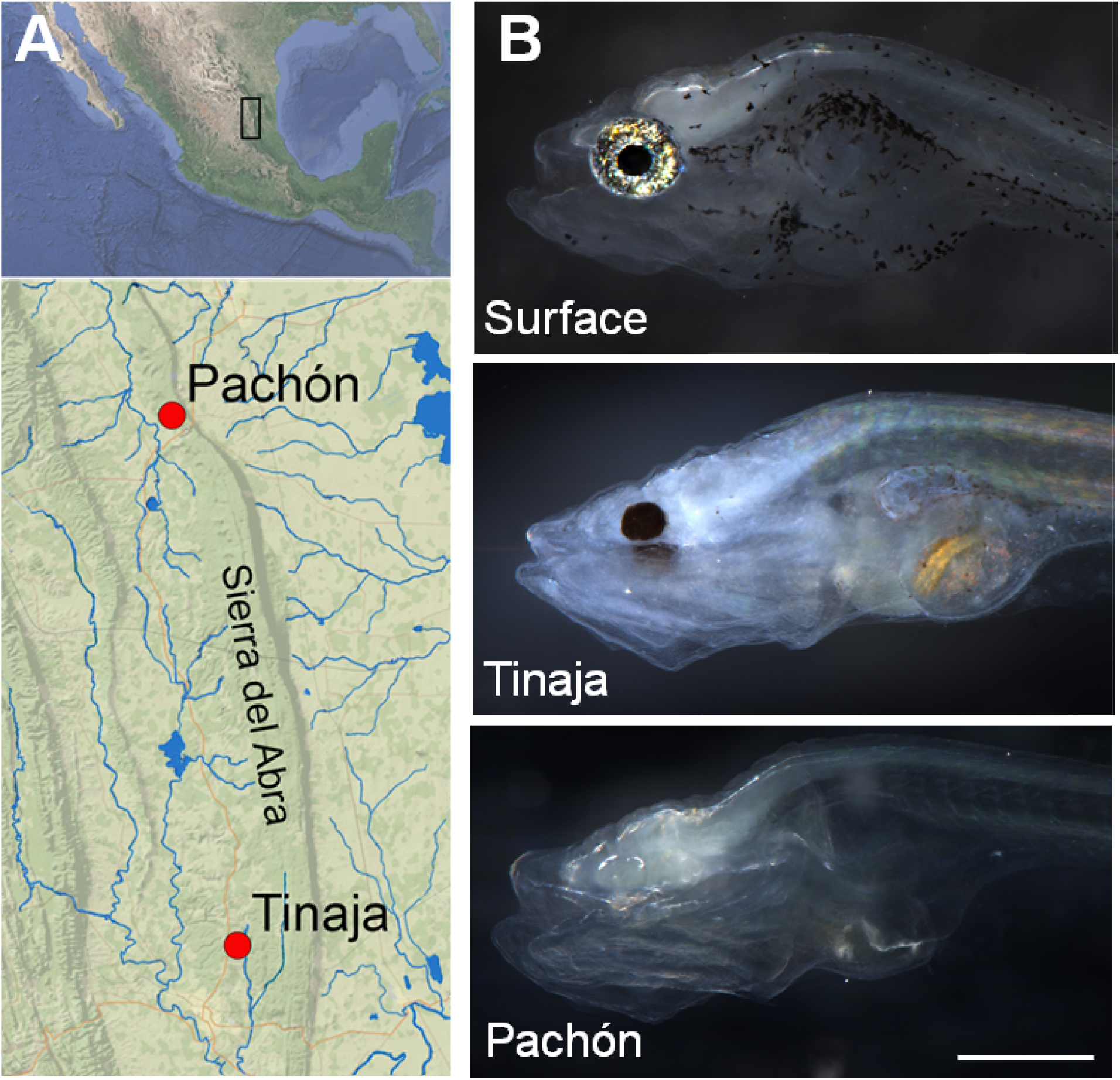
Distribution and morphology of *Astyanax mexicanus* morphs used in this study. A) Tinaja and Pachón morphs are geographically isolated in caves with these names in the Sierra de el Abra region of Mexico (Tinaja: 22°04’34”N 98°58’40”W; Pachón: 22°36’25”N 99°02’55”W). Blue lines represent rivers in the area inhabited by surface morph. Images from Google Earth. (B) Surface fish have pigmented eyes and skin, whereas Tinaja cavefish only have pigmented pupils and Pachón cavefish have no pigment on their skin or eyes, 7dpf. Common scale bar: A. 10 km B. 0.5 mm.

Although adult *Astyanax* cavefish lack eyes, eye development is nevertheless initiated during embryogenesis (reviewed in Jeffery, 2001 and 2009; Krishnan et al., 2017; Emam et al., 2019). To examine the impact of the loss of visual input on the regulation of escape responses, we compared the morphologies of Mauthner cells in normal and dark-reared surface fish, and two lineages of cavefish. Interestingly, we found that the ventral dendrite that receives visual inputs was missing in both lineages of cavefish. To determine whether the loss of the ventral dendrite is caused by evolutionary or homeostatic mechanisms, we examined the morphologies of Mauthner cells of surface fish raised in complete darkness. The ventral dendrites of dark-reared surface fish are larger and more branched that of those reared in daily light cycle. This result shows that in this case, evolutionary and homeostatic mechanisms operate in opposite directions.

## Materials and Methods

### Animal care

Fish larvae originated from a breeding colony and were kept at 20°C water temperature in either a 12h:12h light/dark cycle (12L:12D, surface fish) or constant darkness (24D, surface and cavefish). Breeding was induced by gradually increasing the water temperature from 20°C to 25°C over two days, and fertilized eggs were collected in meshed containers on the bottom of the tanks, based on the procedures of Dr. William Jeffery laboratory. Animal husbandry and experimentation were covered under the Rutgers University IACUC protocol #201702685. NJIT is under Rutgers Newark IACUC oversight.

### Retrograde labeling of Mauthner cells

We injected dye into Mauthner cell axons in larvae at 4 days post fertilization (dpf) by adapting previous protocols for backfilling reticulospinal neurons in teleosts (Fetcho and O’Malley, 1995; Orger et al., 2008). We used borosilicate capillary glass needles (thick-walled, with internal filaments, Cat 1B150F-4, VWR), pulled with a Flaming/Brown pipette puller (Model P-97, Sutter instruments). Injection needles were filled with 62.5 μg/μl of Dextran-TexasRed™, 3,000 molecular weight (Cat D3328, Invitrogen) or 20ug/μl Dextran-Alexa488, 10,000 molecular weight (Cat D22910, Invitrogen). We broke the needle tips by gently swiping against the sharp open end of a Vannas-Tübingen Spring Scissor (Cat 15003-08, Fine Science Tools) to produce a jagged-edged opening.

Fish larvae were anaesthetized before injection in normal Ringer’s solution (116 mM NaCl, 2.9 mM KCl, 1.8 mM CaCl2, 5.0 mM HEPES, pH 7.2; Westerfield, 2000) containing 0.3% ethyl 3-aminobenzoate methanesulfonate (Tricane; MS-222(Cat E10521, Sigma-Aldrich)). Anesthetized larvae were placed onto an agar-coated 35 mm Petri dish, and the injection needle was inserted laterally into the spine with a micromanipulator (M3301R, World Precision Instruments) under a dissecting light microscope (Olympus MVX10).

We injected caudal to the anal pore, approximately halfway between the anal pore and the end of the tail, and took care to avoid injecting into blood vessels. To eject the dye from the pipette tip, we used a PicoPump (PV820, World Precision Instruments) with no more than 20 psi of house-air pressure. We let larvae recover in 24-well dishes in fresh fish water for at least 24 hours to allow for retrograde filling of reticulospinal cells. We screened larvae at 5dpf, under an Olympus MVX10 stereoscope to confirm labeling of Mauthner cells before fixation.

### Fixation

We transferred 5 dpf larvae to 1.5 ml centrifuge tubes (Eppendorf). All fish water was aspirated and replaced with 1.0 ml 4% paraformaldehyde (Electron Microscopy Sciences, USA) in 1X PBS (Phosphate buffer saline, Fisher Scientific, USA). We fixed larvae overnight at 4°C and washed fixative three times with 1X PBS by aspirating and replacing PBS in a centrifuge tube. Fixed larvae can be stored for up to two weeks in 1X PBS.

### Image Acquisition

We dissected the brains of fixed larvae using custom-made tools. These tools consisted of sharpened tungsten wires attached to glass pipettes. We moved each brain to a drop of glycerol in a custom-made coverslip chamber. Briefly, we attached two 20×20 mm #1.5 coverslips to a 20×40 mm #1 coverslip with a room-temperature-hardening mixture made of equal weights of petroleum-jelly, lanolin, and paraffin (VALAP), creating a vertical channel. We also used

VALAP to seal the top and bottom of the channel. We placed fixed brains in the channel, and another 20×20 #1.5 coverslip on top to seal the channel. To Image whole brains we used a laser scanning confocal microscope (Leica SP8; microscope: DM6000CS objective: Leica 40X HC PL APO oil or Leica 63X HC PL APO oil objective with resolutions of up to 0.35 μm, and 0.32 μm, respectively, laser lines: 561 nm (for Dextran-TexasRed™) and 488 nm (for Dextran-Alexa488)). We collected multiple optical slices (thickness optimized by the confocal software) that contained the complete dendritic structure of both Mauthner cells in each brain.

### Image processing and Measurements

We processed images and measured features using a dedicated software package (LAS X, Leica Microsystems). First, we used the viewer panel to adjust the brightness and contrast of the 3D images. Next, the line tool under the measurement function was used to obtain the total dendritic length, the width of the branch, and the first branching point. Processing of images involved using a white noise reduction filter, applied at levels between 1 and 7, to reduce background noise in the LAS X software. We cropped the images using Adobe Illustrator to visually isolate the Mauthner cells from neighboring reticulospinal neurons labeled in the hindbrain.

### Statistical Analysis

We used one-way ANOVA tests with a post-hoc Tukey test to calculate the p-value and determine if the variances of groups were significantly different (Fig 3A, C, D). We calculated p-values for figure 3B with a Wilcoxon test for nonparametric values. We considered an alpha value of 0.05 to be significant.

## Results and Discussion

### Here, we examined two time scales for changes in the morphology of a dendrite involved in visually guided behavior: early development and evolution

We expected both time scales to follow the same path, and our main findings show that the lack of light decreases the size of the dendrite in an evolutionary time scale (tens of thousands of years), but during development (days) darkness has the opposite effect, increasing the size and complexity of the visual dendrite. A possible outcome was that cavefish maintained their visual dendrites after cave adaptation, and these dendrites were co-opted by another sensory modality. Another reasonable outcome would be that dark reared surface fish lost, or at least decreased, visual dendrites. As our results show, neither outcome was seen in *Astyanax*. Instead, reduced input during the time scale of development resulted in an increase in dendrites, opposite to the evolutionary time scale.

The goal of this work was to understand the relationship between evolutionary change and homeostatic regulation. Conceptually, homeostatic mechanisms hinder changes, whereas evolutionary processes drive them. Here, we compared morphological homeostatic regulation to evolutionary changes in larval *Astyanax*. The evolutionary changes were caused by adaptation to troglobitic life, including the loss of eyes and concomitant changes in neural control networks that rely on vision. We examined homeostatic regulation by rearing surface fish in complete darkness, eliminating visual signals. Each of these groups was compared to the basal morphology exhibited by surface fish raised in alternating light and dark epochs like those that the fish experience in nature.

Each Mauthner cell has two primary dendrites: a lateral dendrite (LD) that receives auditory input and a ventral dendrite (VD) that is thought to receive visual input (Medan et al., 2018). We examined Mauthner cells by retrograde labeling with dextran-conjugated dyes in surface larvae raised under one of two conditions: 12L: 12D cycle (control) or 24h dark, to determine if morphological differences may arise in the dendrite if visual inputs were removed. In surface larvae raised under control conditions (Fig. 2C, Supplemental Movie 1) VDs always began with a 2.98±0.5 μm thick proximal section with several finer processes protruding medially (Fig. 3B). The proximal section branched once after 32.86±7.83 μm into two terminal branches (Fig. 3CD). In surface fish raised in constant darkness (Fig. 2D), VD proximal sections were thicker and longer (4.5±0.89 μm, 43.09±9.95 μm, Fig. 3CD) and often arborized into three or more terminal branches (Fig. 3C) and also displayed finer processes propagating off the length of the dendrite. In Tinaja cavefish (Fig. 2E), VD did not branch (Fig. 3B) and appeared thin (diameter 1.79±1.29 μm, Fig. 3C) and truncated with finer processes that protrude medially. Pachón cavefish did not have detectable VDs of any length (Fig. 2F).

**Fig. 2.**
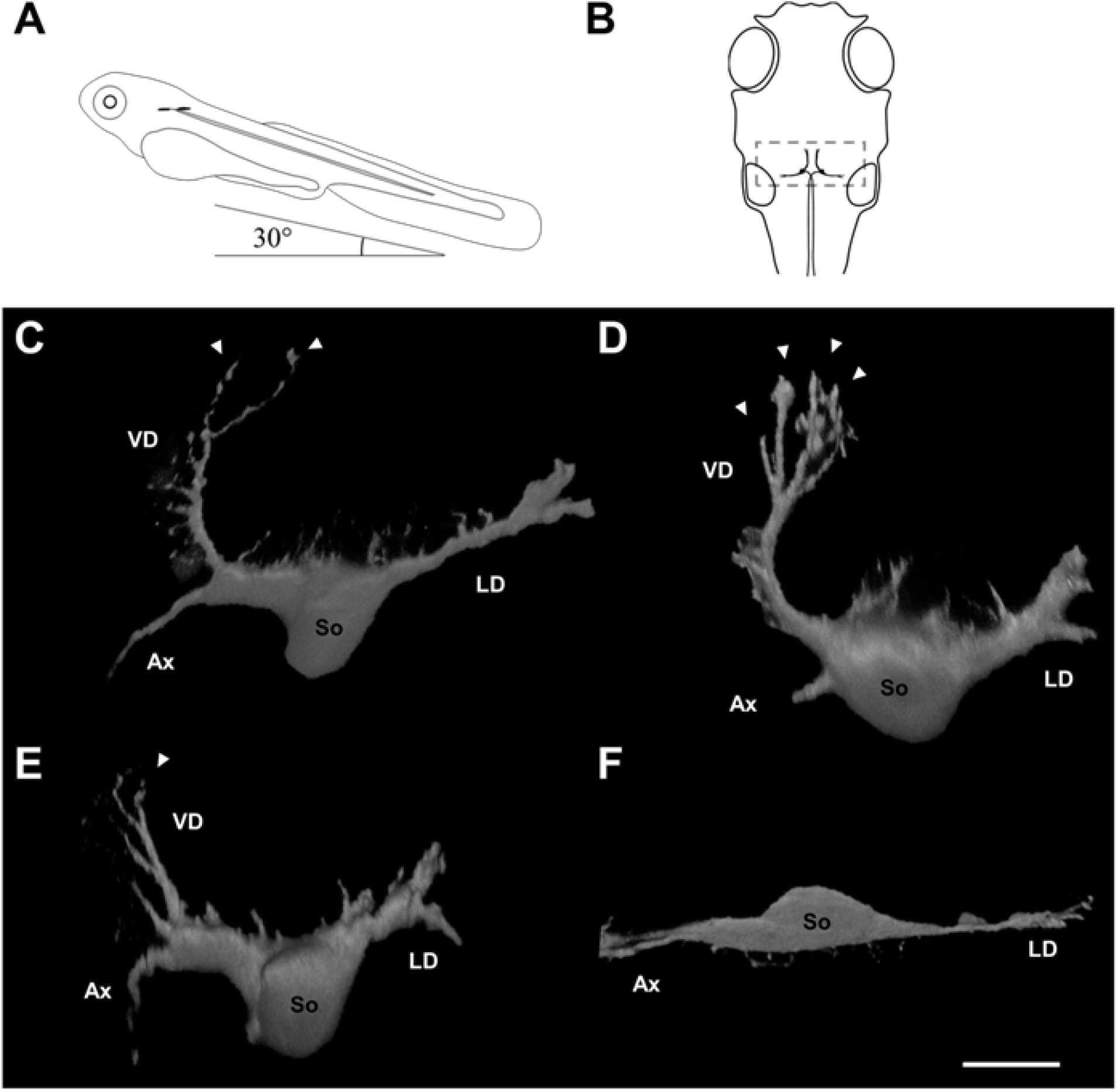
Neuronal morphology differs among Mauthner cells in surface fish raised in the light and dark, and between Tinaja and Pachón cavefish. (A) Larvae were positioned with a nose-up pitch angle of 30° to allow visualization of the ventral dendrite for confocal imaging (lateral view). (B) Position of paired Mauthner cells (black) in the head from a 30-degree pitched dorsal view. Note the dendritic area (gray box) and contralateral axons proceed caudally. (C--F) volume reconstructions of right Mauthner cells. (C) Mauthner cell of surface fish raised under 12L:12D conditions. (D) Mauthner cell of surface fish raised in darkness. Note more dendritic tips on the ventral dendrite. (E) Mauthner cell of Tinaja cavefish. Note the ventral dendrite is smaller. (F) Mauthner cell of Pachón cavefish. Note smaller soma and absent ventral dendrite. VD: ventral dendrite, LD: lateral dendrite, So: soma, Ax: axon, arrowheads: dendritic tips. All neurons from 5 dpf larvae. Scale bar: 20 μm.

The diameter and length of the proximal section, number of dendritic tips, and overall length of VDs in control-raised surface fish were significantly different from those in other morphs and raising conditions (Fig. 3). The two cavefish subtypes were also significantly different from each other in those measurements. VDs of surface fish larvae raised in constant darkness were significantly longer (84.35±11.46 μm, p=0.0009), while VDs of Tinaja larvae were significantly shorter (22.67±18.36, p=0.0001) than control-raised surface morph (54.76±8.67, Fig. 3A). Since Pachón cavefish larvae did not have VDs, their length was recorded as 0 (p<0.0001 to control raised surface fish and p=0.00533 to Tinaja cavefish; Fig. 3A).

**Fig. 3.**
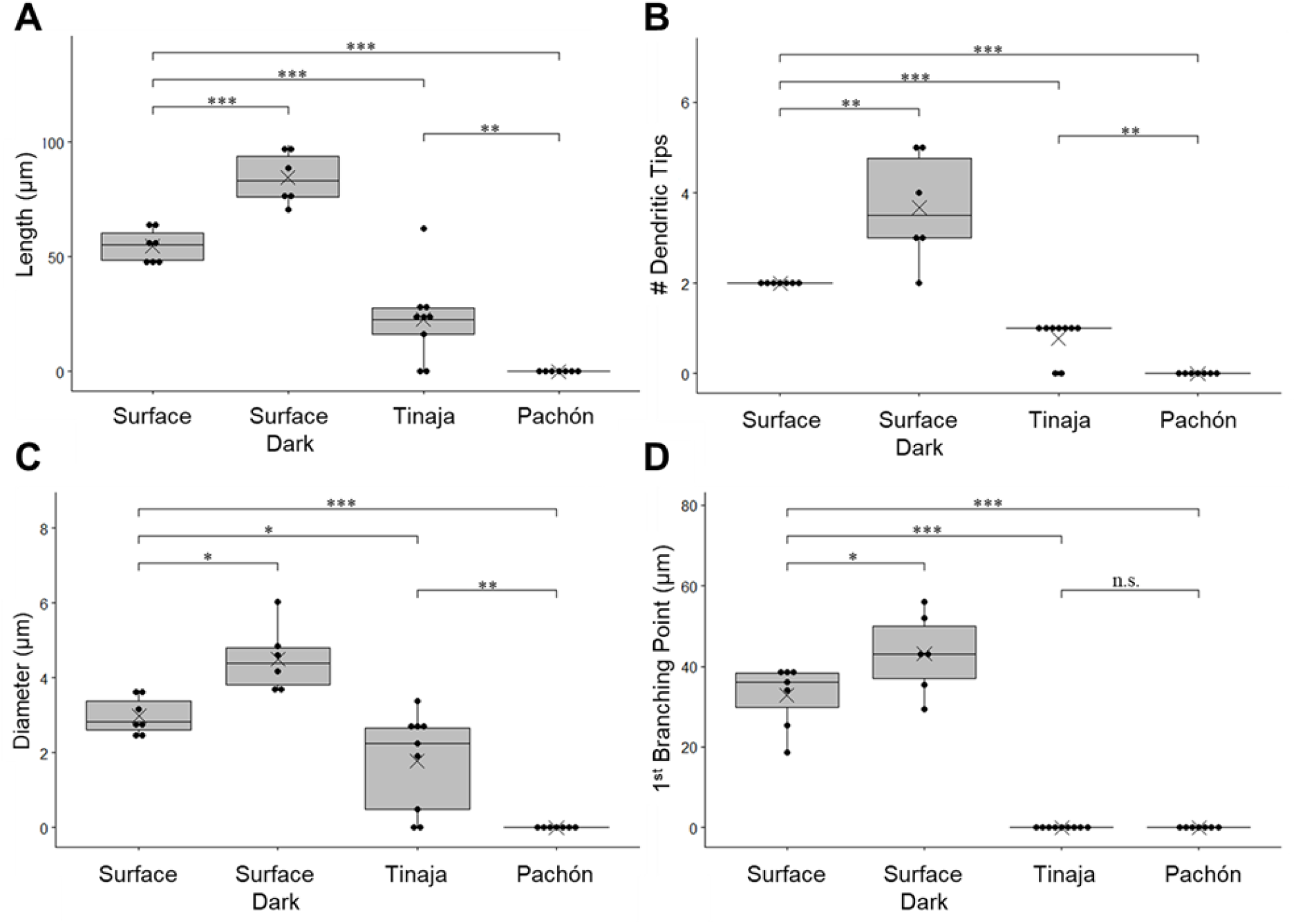
Mauthner cell ventral dendrites of surface fish increase in length and branching when raised in darkness, while cavefish dendrites are smaller or absent. (A) Total dendritic length. (B) Number of dendritic tips (C) Ventral dendrite diameter at its thickest position (D) Distance of first branching point from the cell body. Box plots: demonstrates the distribution of data by showing upper and lower quartiles. dots: data points, line across box: median, X: mean, *** p ⋜ 0.001, **p⋜ 0.01,* p ⋜ 0.05, n.s. > 0.05. The numbers of neurons (animals) are Surface 12L:12D: n= 7 (5), Surface 24D: n= 6 (3), Tinaja: n= 9 (5), Pachón: n= 7 (4).

The number of dendritic tips is a measure of terminal branching (Fig.3C). VDs of surface fish larvae raised in constant darkness were more arborized than control-raised surface morph (p=0.0055), while VDs of Tinaja larvae were less arborized because they always had a single dendritic tip (p=0.00033). Since Pachón cavefish larvae did not have VDs, their numbers of dendritic tips were recorded as 0 (p=0.00041 to control raised surface fish and p=0.0032 to Tinaja cavefish; Fig. 3C). The proximal section of VDs of surface fish larvae raised in constant darkness was thicker (p=0.01936) and longer (p=0.02) than control-raised surface morph (Fig.3BD). Because the VDs of Tinaja did not branch and were absent in Pachón cavefish larvae, there was no meaning to a proximal section, and the distance to the first branching point was recorded as 0 (both cavefish subtypes p<0.0001 to control raised surface fish Fig. 3D).

One of the most intriguing questions in Biology is how evolution changes phenotypes and how evolutionary pressure interacts with homeostatic changes that occur on a much shorter time scale. Do homeostatic responses provide a mechanism for evolutionary modifications? Or conversely, are evolutionary modifications the outcomes of generations of unbalanced homeostatic responses? The latter seems to be a simple and straightforward explanation of how adaptation could be influenced by homeostatic mechanisms. However, here we have found at least one example of a homeostatic response that leads to the opposite outcome than an evolutionary adaptation.

We have shown that *Astyanax mexicanus* surface fish display the commonly found Mauthner cell morphology, which is shared among fishes and amphibians. However, the cavefish morphs have either much smaller or absent ventral dendrites, depending on the duration of the cavefish lineage isolation in the darkness of a cave (Fig. 2).

To examine the influence of the visual environment in the developing circuit, we raised surface fish larvae in constant darkness. We predicted that ontogeny would recapitulate phylogeny, but instead in the surface fish that were raised in the dark, the dendrite that is innervated by the visual system was not only longer, but also much more branched (Fig. 2). It has been a long-standing question why in rivers with so many species of teleosts, only *Astyanax* has been the one to colonize caves successfully, and multiple times (Bilandžija et al., 2020; Yoffe et al. 2020). Here, we added neuromorphological homeostasis to the body of evidence that argues for *Astyanax* as a particularly plastic species, with advantages in dispersal. Consequently, homeostatic mechanisms are under natural selection that provide adaptation to constant darkness.

## Statements

All papers must contain the following statements after the main body of the text and before the reference list:

## Acknowledgement

We would like to thank Marina Yoffe for the collection of *Astyanax* eggs and Dr. Eric Fortune for their helpful comments on the manuscript. We thank Kathryn Gallman for the fish care.

## Statement of Ethics

No humans were used in this study.

## Funding Sources

Grant support: NIH R15EY027112.

The data that support the findings of this study are available from the corresponding author upon reasonable request.

## Author Contributions

ZT, DS, KS, and GH conceived the project. ZT, DS, and DR collected and analyzed the data. All the authors discussed the project throughout and wrote the paper.

